# The avian W chromosome is a refugium for endogenous retroviruses with likely effects on female-biased mutational load and genetic incompatibilities

**DOI:** 10.1101/2020.07.31.230854

**Authors:** Valentina Peona, Octavio M. Palacios-Gimenez, Julie Blommaert, Jing Liu, Tri Haryoko, Knud A. Jønsson, Martin Irestedt, Qi Zhou, Patric Jern, Alexander Suh

## Abstract

It is a broadly observed pattern that the non-recombining regions of sex-limited chromosomes (Y and W) accumulate more repeats than the rest of the genome, even in species like birds with a low genome-wide repeat content. Here we show that in birds with highly heteromorphic sex chromosomes, the W chromosome has a transposable element (TE) density of >55% compared to the genome-wide density of <10%, and contains over half of all full-length (thus potentially active) endogenous retroviruses (ERVs) of the entire genome. Using RNA-seq and protein mass spectrometry data, we were able to detect signatures of female-specific ERV expression. We hypothesise that the avian W chromosome acts as a refugium for active ERVs, likely leading to female-biased mutational load that may influence female physiology similar to the “toxic-Y” effect in *Drosophila*. Furthermore, Haldane’s rule predicts that the heterogametic sex has reduced fertility in hybrids. We propose that the excess of W-linked active ERVs over the rest of the genome may be an additional explanatory variable for Haldane’s rule, with consequences for genetic incompatibilities between species through TE/repressor mismatches in hybrids. Together, our results suggest that the sequence content of female-specific W chromosomes can have effects far beyond sex determination and gene dosage.

## Introduction

Many organisms exhibit a genetic sex determination system where a pair of sex chromosomes guides sex development [1]. There are two major genetic sex determining systems: the XY system with male heterogamety (XX females and XY males) and the ZW system with female heterogamety (ZW females and ZZ males), whereby the Y and W are the sex-limited chromosomes (SLCs).

Sex chromosomes generally evolve from a pair of autosomes [2] that acquire a sex-determining locus and locally suppressed recombination around that locus [3,4]. The non-recombining region may remain very small, keeping the two sex chromosomes largely homomorphic. Conversely, in heteromorphic sex chromosomes the non-recombining region may expand over time until only a small pseudo-autosomal region (PAR) remains recombining, while the rest of the SLC diverges, degenerates or loses genes, and accumulates repeats [5]. The evolution of the non-recombining region of the SLC is mostly shaped by its low recombination rate. Its associated low effective population size drastically decreases the efficacy of selection [6] (i.e., accentuating the effects of drift and linked selection) and makes these chromosomes vulnerable to the accumulation of slightly deleterious mutations (e.g., through Muller’s ratchet and Hill-Robertson interference mechanisms), such as repeats [3,7].

Because of their low gene content and high repeat density, SLCs were thought to not have any effect beyond sex determination and gonadal development, remaining largely understudied or even absent in the majority of the genome assemblies and studies [8]. However, recent studies on SLCs, especially in humans and other model organisms, have shown that they play roles in human diseases [9,10], male infertility [11], determining sex-specific traits [12], shaping the genome-wide heterochromatic landscape [13], exerting epistatic effects [14–16], reproductive isolation [17], and suppressing meiotic drivers on other chromosomes (e.g., through RNAi pathways) [18].

While Y chromosomes of mammals and flies have recently received considerable attention, the evolutionary implications of W chromosomes in any organism are still poorly understood. Here we provide the first evidence that the avian W chromosome is not merely a graveyard of repetitive elements but a refugium of potentially active transposable elements (TEs) that likely have sex-specific implications. Bird genomes are known to be repeat-poor with a mean TE content of <10% [19], but the first female assemblies based on short [20] or long reads [21–23] showed that the non-recombining W chromosome is >50% repetitive and especially rich in endogenous retroviruses (ERVs). By analysing reference-quality genomes of six species spanning the avian Tree of Life from both Paleognathae (emu with homomorphic sex chromosomes) and Neognathae (chicken, Anna’s hummingbird, k•k•p•, paradise crow, zebra finch with heteromorphic sex chromosomes), we demonstrate that the avian W has generally accumulated ERVs and likely contains active ERVs as indicated by signatures of transcription and translation of W-linked ERVs. We therefore hypothesise that the W is a sex-specific source of genome-wide retrotransposition and genome instability, with the male/female difference in ERVs dictating the degree of repercussions on sex differences in physiology and reproductive isolation.

## Results and discussion

### Enrichment of ERVs on the W chromosome

We analysed six avian genomes spanning the avian Tree of Life (**Figure 1A**) and representing the current standard for reference-quality genome assemblies [23,24]. Autosomes had between 6 and 12% TEs on average (**Figure 1B**, **Supplementary Table S2**, **Supplementary File S1**) and the Z chromosome had similar or slightly more TE densities (5-17%), while the W chromosome stood out as having ~22-80% TEs (**Supplementary Table S2**). Notably, we also found the homomorphic W chromosome of emu to be richer in TEs than the autosomes and Z (22% vs. 6.4% and 5.6%). Generally, the Z chromosome exhibited a TE landscape more similar to the autosomes than to the W chromosome, both regarding abundances and types of TEs (**Figure 1B**, **Supplementary Table S2**). While long interspersed elements (LINEs) from the Chicken Repeat 1 (CR1) superfamily were the dominant repeats on autosomes and Z (cf. [19,25]), ERVs were the major component of the W chromosome and accounted for more than 50% of the assembled chromosome itself (**Supplementary Table S2**).

**Figure 1.**
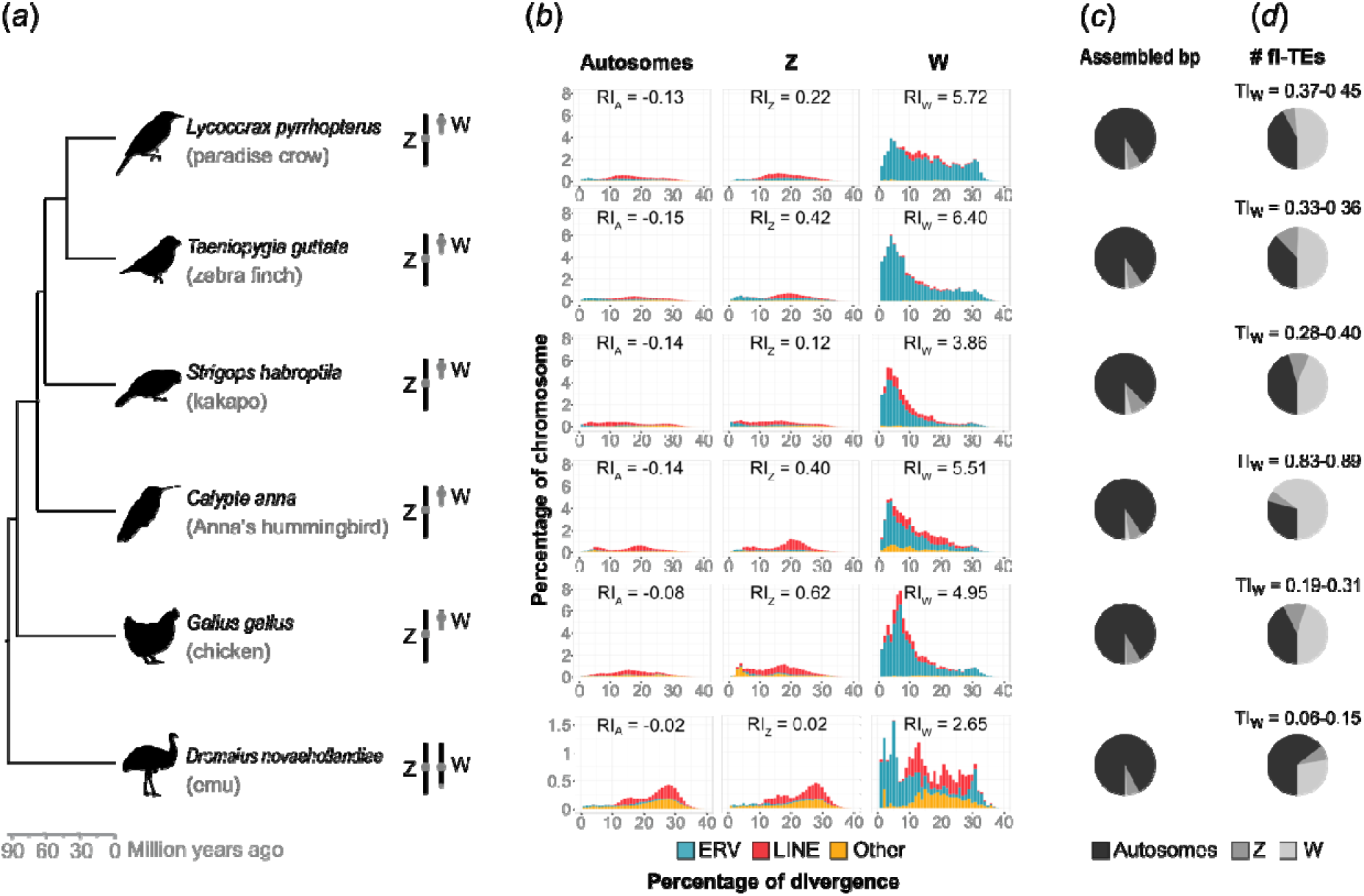
Massive accumulation of endogenous retroviruses (ERVs) on W chromosomes of six female reference-quality genome assemblies spanning the avian Tree of Life. *(a)* Avian time tree after [34] with schematic homomorphic or heteromorphic sex chromosomes [32]. *(b)* TE landscapes of autosomes and sex chromosomes as stacked bar plots. Abundance of interspersed repeats (bp occupied) normalised by chromosome size plotted against percentage of divergence calculated as Kimura 2-parameter distance to consensus. The refugium index (RI) for interspersed repeats on autosomes, Z and W is indicated for each species. *(c)* Comparison of autosome and sex chromosome assembly sizes as pie charts. *(d)* Comparison of full-length TE numbers (mainly ERVs) on autosomes and sex chromosomes as pie charts. The toxicity index (TI) of the W chromosome is reported for each species as the range between estimates from RetroTector or LTRharvest + LTRdigest (**Table 1**).

ERVs are Long Terminal Repeat (LTR) retrotransposons deriving from germline-inherited retrovirus integrations and exist mainly in two genomic forms [26,27]: 1) full-length elements with two long terminal repeats (likewise called LTRs) flanking its protein-coding genes necessary for retrotransposition; 2) solo-LTRs resulting from homologous recombination between the two flanking LTRs. Only full-length elements are capable of autonomous retrotransposition. Using RetroTector and LTRharvest/ LTRdigest [28–30], we annotated full-length ERVs (fl-ERVs; **Supplementary Files S2-3**) and detected a large proportion of fl-ERVs on the W chromosome compared to the rest of the genome (**Figure 1C**, **Table 1**, **Supplementary Tables S3-12**). Despite the fact that the W chromosome accounted only for the 1-3% of the total length of assembled chromosomes (**Figure 1C**, **Supplementary Table S3**), this chromosome carried the same or higher numbers of fl-ERVs than the autosomes altogether, with the exception of emu with half the number on W than autosomes together (**Figure 1D**, **Table 1**). The distribution of fl-ERVs deviated significantly (χ^2^ test, <u>P</u>-values < 0.01) from a random distribution across all chromosomes (**Supplementary Table S4**), with an impoverishment of total ERV-derived bp on the autosomes (0-0.4 times fewer bp than expected) and an extreme accumulation on the W (12-54 times more bp than expected; **Supplementary Table S12**). In contrast, we identified a negligible amount of other full-length TEs per genome (0-11 DNA transposons, 0-8 CR1 LINEs; **Supplementary Table S4**).

**Table 1.**
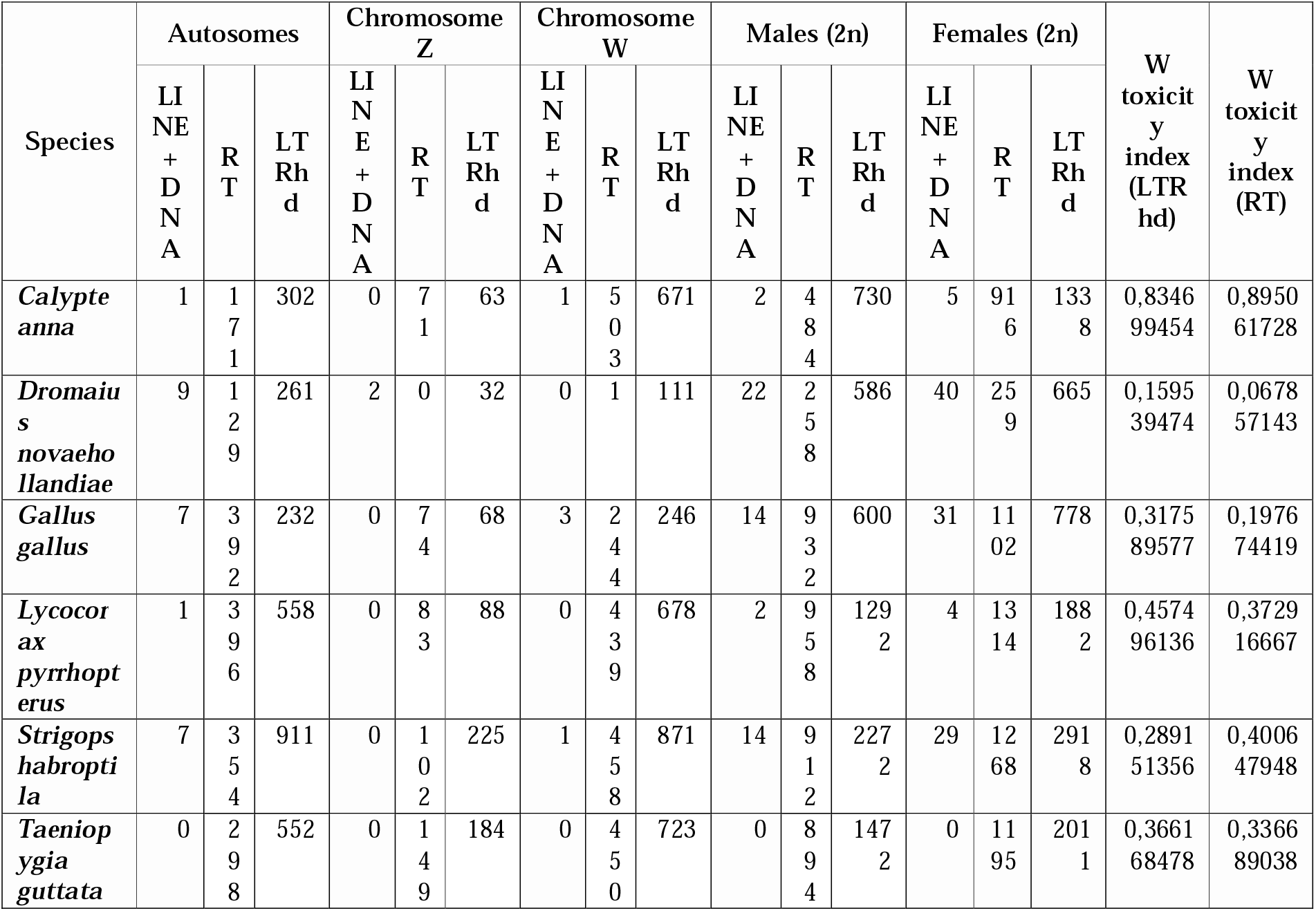
Number of full-length endogenous retroviruses (fl-ERVs) and other full-length TEs (LINE + DNA) found on autosomes and sex chromosomes. fl-ERVs were identified using either RetroTector (RT) or LTRharvest + LTRdigest (LTRhd).

We propose a “refugium index” (formula **1.1**) to quantify the excess accumulation of TE-derived bp on an SLC relative to the rest of the genome by comparing the observed and expected abundance of TEs. Positive values of the refugium index indicate an excess of TEs while negative values a depletion of TEs. Since only a subset of TE copies are usually capable of (retro)transposition, we propose a “toxicity index” as a quantitative measure for the excess of intact TE copies in the heterogametic vs. homogametic sex through the presence of an SLC (formula **1.2**). The excess is calculated by comparing the number of full-length TEs in the diploid state in the two sexes. The toxicity index indicates a non-toxic SLC when equal to 0, toxicity of SLCs when positive, and toxicity of Z or X when negative. The term “toxicity” pays tribute to the recently proposed “toxic Y” hypothesis in *Drosophila* [13], which suggested that an excess of Y-specific active TEs can lead to male-biased transposition and genome instability, together likely detrimental to the genome and the organism. For birds, we calculated the toxicity index as the excess of fl-ERVs carried by diploid females compared to diploid males (**Table 1**), suggesting that females with heteromorphic sex chromosomes carried between 20 to 90% more fl-ERVs than males, and that even the emu has 7-16% more fl-ERVs in females than males despite largely homomorphic sex chromosomes [31,32]. We assume this phenomenon to reflect that the non-recombining region of the W, no matter how big or small, constantly accumulates large quantities of new TEs. It is important to note that, given the difficulties in assembling SLCs even with long-read sequencing technologies [8,23,24,33], the W chromosome models are likely to be less complete than the other chromosomes. We thus consider our W repeat annotations as well as indexes to be conservative estimates for the true repeat content.

Our results suggest that the avian W chromosome is acting as a refugium for intact and thus potentially active TEs, particularly ERVs, which may have numerous implications. We thus propose the “refugium hypothesis” for SLCs in general: the accumulation of TEs on the SLC leads to an excess of intact TEs in the heterogametic sex, with a toxic effect absent from the SLC-lacking homogametic sex. This sex-specific toxic effect may manifest itself as sex-biased mutational load, genomic instability, ageing, and genetic incompatibilities as the result of SLC-linked TE activity and heterochromatin dynamics (explained below). To quantify and test the refugium hypothesis in any sex chromosome system of interest, we introduced two indexes above: the refugium index to measure the density of TE-derived bp on the SLC relative to the remaining chromosomes; and the toxicity index to measure the number of intact TEs (i.e., full-length copies of LTRs, LINEs and DNA transposons) in the heterogametic sex relative to the other sex.

### Transcription and translation of W-linked ERVs

Considering the exceptionally high number of W-linked fl-ERVs, we tested whether the avian W chromosome harbours a potentially active load of ERVs specific to females. In the absence of available retrotransposition assays for birds, we regarded the transcription and translation of W-linked ERVs as proxies of their activity. We identified W-linked single-nucleotide variants (SNVs) within ERVs by mapping genome re-sequencing data from male and female individuals, as well as female transcriptome data, to consensus sequences of our repeat library (**Supplementary File S5**). We consider this to be a conservative subset of W-linked SNVs because we required each SNV to be present in all females and absent in all males per species. However, the paradise crow dataset that contain only one male likely gave rise to false positive W-linked SNVs. We then traced the presence of ERV proteins in the male and female proteome data available for white leghorn chicken.

We analysed zebra finch, paradise crow, chicken, and emu for W-linked SNVs in genome re-sequencing and RNA-seq data mapped against ERV consensus sequences. In each species, we found between 52 and 332 ERV subfamilies with W-linked SNVs (**Table 2**), with ERVL subfamilies being the most represented (**Supplementary Table S13**) and found evidence for the transcription of between 12 and 182 ERV subfamilies in female gonads or female pectoral muscle (**Table 2**, **Supplementary Table S13**). Our estimates of transcribed W-linked ERVs are likely just the tip of the iceberg, because we expect to identify W-linked SNVs only if those ERVs did not yet spread in the genome (e.g., very recent variants) or if they accumulated exclusively on the W chromosome (e.g., fl-ERVs only existing as solo-LTRs on other chromosomes). Alongside ERVs, we also identified W-linked SNVs in CR1 LINEs and DNA transposons (**Supplementary Table S13**). Although there is evidence for their transcription, their scarcity of full-length elements makes CR1 LINEs and DNA transposons an unlikely source of mutational load for females.

**Table 2.** Number of female-specific and thus W-linked SNVs detected at the genomic and transcriptomic levels for each species. More details about SNVs in ERVs, LINEs, and DNA transposons are in **Supplementary Table S13**.

| Species | # W-linked SNVs in ERVs | # ERV subfamilies with W-linked SNVs | # transcribed SNVs in ERVs | # transcribed ERV subfamilies |
| --- | --- | --- | --- | --- |
| *Dromaius novaehollandiae* | 764 | 58 | 82 | 12 |
| *Gallus gallus* (red junglefowl) | 2088 | 52 | 671 | 28 |
| *Gallus gallus* (white leghorn) | 6385 | 166 | 3534 | 102 |
| *Lycocorax pyrrhopterus* | 1591 | 198 | 42 | 21 |
| *Taeniopygia guttata* | 15012 | 332 | 3306 | 182 |

Next, we analysed the overall RNA expression level of ERVs in male and female gonads of emu, chicken, and zebra finch via RNA-seq read mapping to genomic regions annotated as LTR or ERV fragments by RepeatMasker (**Figure 2a**). Overall, females expressed such ERVs more highly than males, with the single Z chromosome of females showing expression levels that matched the two male Z chromosomes. This pattern contrasts with incomplete dosage compensation of Z-linked genes in birds indicated by usual 2-fold higher expression level of Z-linked genes in males [35,36]. Assuming that some of these ERVs are full-length and capable of retrotransposition, female gonads would thus be exposed to a greater mutational load. Furthermore, many of the autosomal and Z-linked ERVs showed differentially expression towards females (**Supplementary Figure S1**).

**Figure 2.**
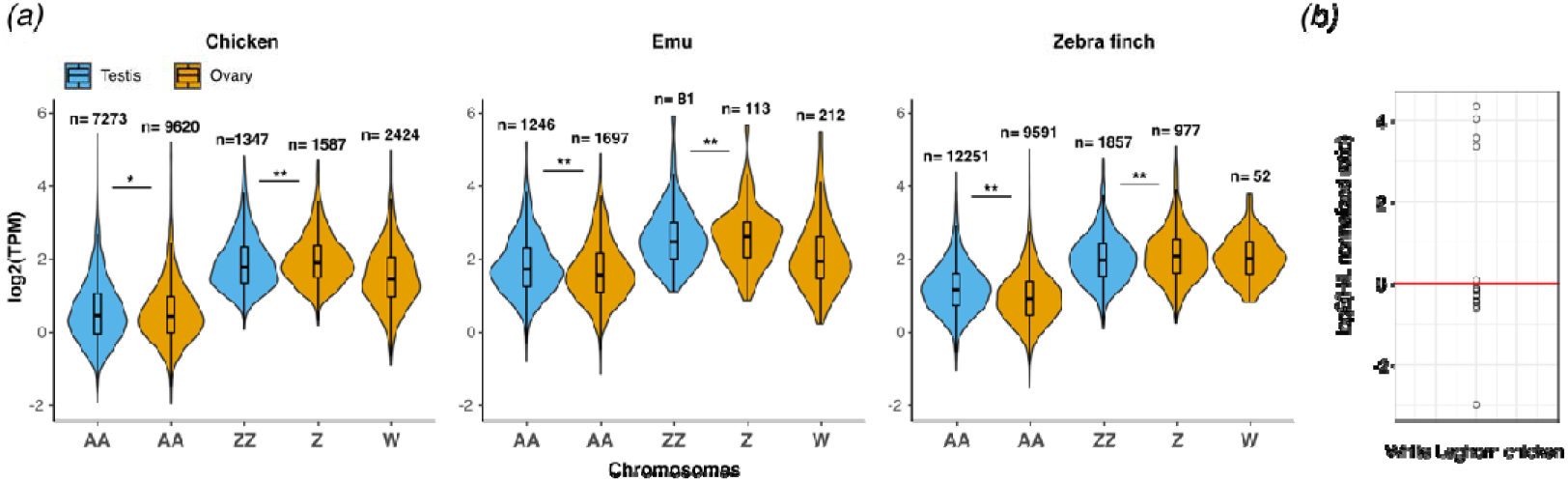
RNA and protein expression of ERVs in male testes and female ovaries of different birds. *(a)* RNA expression of all genomic copies/fragments (n) annotated as LTR or ERV by RepeatMasker. Violin plots show ERV expression levels by chromosomes using the average number of RNA-seq reads across replicates, normalized for ERVs length and library size mapping to each chromosome (A, Z, and W) from ovaries (blue) and testes (orange). Significance values calculated using the Wilcoxon test (*P-value < 0.05, **P-value < 0.01). TPM: Transcripts Per Million reads. *(b)* Scatterplot of log2 fold-change of the H/L SILAC ratio of ERV-related peptides from 15 ERV subfamilies expressed in chicken ovaries (H) and testis (L). Values above 0 indicate proteins with female-biased expression while values below 0 are proteins with male-biased expression. The ratios for 5 translated LINE-related peptides are found in **Supplementary Files S6**. H: heavy protein labelling; L: light protein labelling.

Finally, we analysed protein mass spectrometry data of white leghorn chicken gonads [37] with MaxQuant [38] for the presence of TE-related proteins and found a higher quantity of more of these proteins expressed in females than in males as indicated by a high H/L SILAC ratio (**Figure 2b**, **Supplementary File S6**). Together, these results demonstrate that some W-linked ERVs are transcribed and that females have more ERV translation than males, and that W chromosomes thus feature fl-ERVs potentially able to retrotranspose. Given our present data, we cannot distinguish whether this higher ERV translation stems solely from the W chromosome but it is plausible that the presence of a SLC causes a higher TE activity (similarly to what happens in *Drosophila* [13]).

### Sex-biased implications for mutational load

SLCs have been largely considered inert chromosomes with few effects beyond sex determination and gonadal development because of their low gene content (e.g., only 13 genes on Drosophila Y [39] and 28 genes on chicken W [21]). However, accumulating evidence shows that SLCs can have additional effects [12,40,41]. For example, it is important to highlight that the Y-linked regulatory variation within populations of *Drosophila* can have genome-wide epistatic effects [14–16,42]. This Y-linked regulatory variation cannot be explained simply by regulatory variation of the protein-coding genes and it has been proposed that the variability in Y repetitive content and structural variation are responsible for re-shaping the genome-wide heterochromatin landscape [43]. This hypothesis is known as the heterochromatin sink model, suggesting that large heterochromatin blocks on SLCs act as a sink for the heterochromatin machinery and thereby reduce the efficiency of heterochromatin maintenance elsewhere relative to the SLC-lacking sex [13,43].

Recently, the Y chromosome repeat content has been linked to the destabilisation and loss of heterochromatin, which in turn is correlated to the shorter lifespan of the heterogametic sex [13,44]. By using *Drosophila melanogaster* experimental lines with different Y dosages (XO males, XXY females, XYY males), Brown et al. [13] showed that the presence and number of Y chromosomes carried are correlated with shorter lifespans. It was thus suggested that the Y itself is “toxic” for the entire genome and organism, and this toxicity is caused by the Y-linked load of active TEs [13,45,46] whose expression is unleashed by heterochromatin loss. Possibly, the dysregulation of TEs due to heterochromatin loss is also associated with laminopathic diseases in *Drosophila* and humans [47]. According to the refugium hypothesis proposed here, we predict that in species with a high toxicity index (i.e., excess of intact TEs on the SLCs and/or paucity thereof in the rest of the genome) this toxic effect will be more accentuated (**Figure 3A** and **B**). The toxic-Y hypothesis has been recently investigated from a theoretical point of view in vertebrates which proposes that the expression of recessive mutations on X/Z chromosomes is the cause of the shorter lifespan in the heterogametic sex. It is important to note that reduced female lifespan in birds has been document in many species [52–55]. Sultanova et al. [48] used the sizes of Y and W relative to X and Z as a proxy for toxicity, i.e., assuming that smaller SLCs are more repetitive. Although the correlation between the Y size and relative lifespan in mammals was strong, the authors did not find such a correlation for the W in birds. We note that while SLC size relative to X/Z size might indeed correlate negatively with the overall repeat content (i.e., satellites and fragmented TEs), this might not necessarily be informative for the number of intact TEs. Therefore, we propose that our toxicity index could be a more suitable proxy for toxicity since it considers the sex differences in the load of intact and (potentially) active TEs. Among the six birds compared here, emu and Anna’s hummingbird would be those with the lowest and highest toxicity indexes, and it remains to be tested if this indeed is a better predictor of female lifespans.

**Figure 3.**
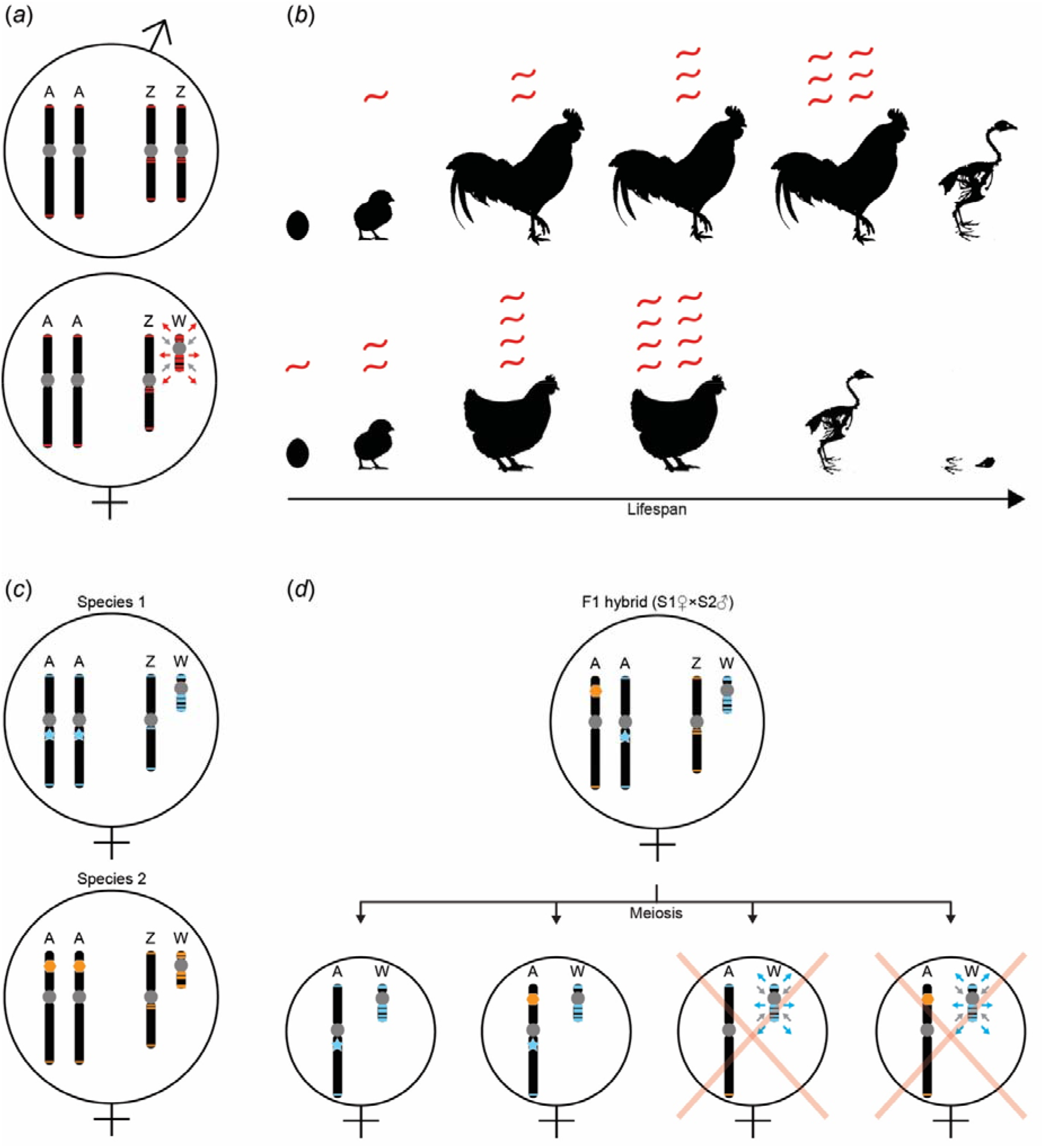
Synthesis of the consequences of the refugium hypothesis on micro- (*a* and *b*) and macroevolutionary (*c* and *d*) time scales. For simplicity, a schematic example of avian sex chromosomes is shown, but we expect these consequences for any ZW or XY system with SLC-linked intact transposable elements (TEs). *(a)* Simplified karyotypes of male and female birds indicating that the W chromosome is a TE refugium and heterochromatin sink. Grey circles: centromeres and heterochromatin blocks; red lines: intact TEs; red arrows: TEs spreading away from the W chromosome; grey arrows: heterochromatin deposition on the W chromosome. *(b)* The “toxic” effect of the gradual de-repression of TEs during an organism’s lifetime is more accentuated in females carrying more intact TEs than males. The W-linked activity of intact TEs could explain the shorter lifespan of the heterogametic sex. The toxicity of active TEs is represented by the increasing number of transcripts in red as a proxy for genome-wide TE insertions. *(c)* Simplified karyotypes of two bird species with species-specific TEs (blue and orange lines) and sequence-specific TE repressors (blue star and orange hexagon). Assuming a rapid accumulation and sequence turnover of TEs especially on the W, diverging species or populations may quickly acquire different TE/repressor repertoires. *(d)* Genetic incompatibility due to the W chromosome in a female F1 hybrid between species 1 and species 2 of panel C. Schematic example of four possible meiotic products (oocytes) of the F1 hybrid, two of which lack the blue repressor of blue TEs because of meiotic recombination between the autosomes. The TE/repressor mismatch may lead to de-repression of W-linked TEs in gametes or embryos and thereby a female-biased reduction in hybrid fitness.

### Sex-biased implications for genetic incompatibilities

In addition to TE mutational load and heterochromatin maintenance influencing organismal physiology, SLC-linked TEs can also play an important role during hybridisation. This point may be not overly surprising in the context of Haldane’s rule, which states that upon hybridisation, if there is a sterile or inviable sex, it will be the heterogametic one. Accumulating evidence suggests that hybrid genome stability can be compromised during mitosis and meiosis by species-specific differences in heterochromatin landscapes leading to uncontrolled TE activity (reviewed by [56]). Furthermore, species-specific families of repeats can induce lagging chromatin at cell division during early embryogenesis (when heterochromatin is first established), leading to chromosome mis-segregation and F2 hybrid embryo death [57]. In the context of the refugium hypothesis, it is important to consider that new and active TEs are one of the main targets of heterochromatinization [58,59], and SLCs could be a source for both sex-specific and species-specific heterochromatin differences.

TEs generally evolve very rapidly in their sequence and usually only few elements remain intact and capable of transposition [60]. In addition, many TE repressor systems are in a sequence-specific arms race (e.g., piRNAs or KRAB-zinc finger proteins [58,61,62]), therefore TE sequences and their repressors can both diverge rapidly between populations and species. Because SLCs rapidly evolve and accumulate repeats [5,18], SLCs are likely sex-specific refugia of species-specific active TEs. Hybrid incompatibility due to TE/repressor mismatches can arise when new TE families are introduced into a naive genomic background (lacking specific repressors), which can lead to the uncontrolled proliferation of such TEs, followed by gene disruption, genome instability [63], and hybrid dysgenesis [17]. TE/repressor mismatches can already occur during meiosis in the F1 hybrids, when recombination can separate the repressor from the controlled TEs (**Figure 3C** and **D**) [64]. Although this scenario can occur in both sexes, we expect that in species with a high number of intact TEs on the SLCs relative to the rest of the genome (i.e., high toxicity index, highly heteromorphic SLCs), there are more chances for a mismatch between a repressor and intact TEs on the SLCs than for other chromosomes (**Figure 3D**).

For the birds analysed here, the W chromosome is likely the main source for genome-wide new TE insertions because it contains 16-50% of all intact TEs in a diploid female. Furthermore, potential TE/repressor mismatches stemming from the W chromosome would also reinforce the observation of reduced mitochondrial (maternally inherited as the W) introgression during hybridisation in birds [65]. Mitonuclear incompatibilities, i.e., mismatches between mitochondrial and nuclear alleles, play a disproportionate role in both intra-species and hybrid incompatibilities [66–69]. In ZW systems, these mitonuclear incompatibilities may be even more exacerbated because of the co-inheritance with the SLCs, and especially when SLCs feature active repeats. Thus, in addition to the preservation of dosage-sensitive genes [21,70], the W represents a reservoir of many different and intact TEs that, through their potential for de-repression in hybrids, may constitute an additional explanatory variable for Haldane’s rule.

## Conclusions

We suggest that the avian W chromosome, no matter how heteromorphic or homomorphic, is a refugium for TEs and specifically fl-ERVs, some of which are expressed and thus potentially capable of retrotransposition. This pattern should be generalisable for all birds given our broad sampling of Palaeognathae, Galloanserae, and Neoaves. We propose that ERVs are continuously shaping W evolution and are one of the major contributors of structural changes of this chromosome. If so, it is reasonable to speculate that ERVs have played a relevant role in the expansion of the non-recombining region of the W (cf. [71]), for example by contributing to the heterochromatinization of euchromatic regions through new ERV insertions.

We hope that the refugium and toxicity indexes proposed here will help testing these hypotheses in avian W chromosomes, and SLCs in general. The toxicity index measures the excess of intact TEs on an SLC, which represents the potential for genome-wide sex-specific mutational load as well as sex-specific genome instability. On the short time scale of individuals, a high toxicity index could lead to larger physiological differences between the two sexes [13]. In the long term, e.g., between populations and species, the accumulation of TEs as measured by the refugium index can have effects on reproductive isolation through TE/repressor mismatches, similarly to the situation in *Drosophila* [17,57]. It is important to underline that the toxicity of SLCs should be linked to the number of intact TEs rather than to the general repetitiveness of the chromosome. Furthermore, the refugium and toxicity indexes can be useful to predict and test hybrid incompatibilities, in addition to measuring the genetic distance between nuclear and mitochondrial genes [72]. We predict that with the increasing availability of genome assemblies based on long reads, these indexes will find applicability across SLCs in general. For birds and their W chromosomes, the possible toxic effect of the W on lifespan requires additional tests *in-vivo* that exclude the effects of the phenotypic sex (e.g., developing systems similar to the four core genotypes in mice [73] or the attached-X/attached-X-Y karyotypes in *Drosophila* [74,75]) and account for confounding ecological factors (e.g., intense sexual competition and predations especially of males).

To conclude, SLCs are not merely refugia for repeats with usually neutral or slightly deleterious effects on SLCs themselves, but SLC-linked intact TEs may have genome-wide effects that could effectively turn SLCs into “toxic wastelands”.

## Materials and methods

### Samples, DNA, RNA and proteome data

We used the female reference-quality genome assemblies of chicken (*Gallus gallus*; GCA_000002315.5; galGal6a), paradise crow (*Lycocorax pyrrhopterus;* accession number pending) [23], emu (*Dromaius novaehollandiae*) [76], Anna’s hummingbird (Calypte anna; GCA_003957555.2; bCalAnn1_v1.p) [24], k•k•p• (Strigops habroptila; GCA_004027225.2; bStrHab1.2.pri) [24] and zebra finch (*Taeniopygia guttata;* GCA_009859065.2; bTaeGut2.pri.v2) [24]. All these six assemblies have chromosome models and we carried out all analyses considering only using assembled chromosomes, i.e., discarding unplaced contigs and scaffolds.

For chicken, Illumina genome re-sequencing libraries were collected for two females and three males of *Gallus gallus gallus* (red junglefowl) from [77] (originally uploaded on NCBI as of undetermined sex) and a female library of *Gallus gallus bankiva* (red junglefowl from Java) from [78]. The sexes of the individuals from [77] were determined using the SEXCMD with default sex markers [79]. Red junglefowl RNA-seq libraries of a female (ovary) and of a male (testes) were retrieved from [80]. We also collected publicly available data for the chicken breed white leghorn, i.e., Illumina genome re-sequencing libraries of one female and three males from [78,81], RNA-seq libraries and protein mass-spectrometry libraries for five ovaries and five testes [37].

For paradise crow, we used one 10X Genomics Chromium linked-read library of DNA from a pectoral muscle sample of a female from [23]. We also newly generated such data for three females and one male using the same methods [23] and generated RNA-seq library from female pectoral muscle (preserved in RNAlater). RNA was extracted with phenol-based phase separation using the TRIzol reagent (ThermoFisher Scientific) following the standard protocol recommended by the supplier, followed by DNase treatment for 30 min using the DNA-free DNA Removal kit (ThermoFisher Scientific). Sequencing libraries were prepared according to the TruSeq stranded total library preparation kit with RiboZero Gold treatment (Illumina, Inc., Cat No.20020598/9). Paired-reads (150 bp) were sequenced on the NovaSeq SP flowcell (Illumina, Inc.).

For zebra finch, we used Illumina genome re-sequencing libraries of four females and four males from [82], and RNA-seq libraries of two ovaries and one testis from [83,84].

Finally, for emu we collected Illumina genome re-sequencing libraries of two females and two males from [85–87], and RNA-seq libraries for seven ovaries and five testes from [86,88].

More details and accession numbers for all the libraries and genomic sequences here utilised can be found in **Supplementary Table S1**.

### Repeat annotation

To best annotate repeats in all six avian species, we made sure to have species-specific repeat predictions for each. The repeat libraries of chicken, paradise crow and zebra finch were already manually curated elsewhere [23,89,90] while species-specific repeat libraries did not exist for emu, Anna’s hummingbird and k•k•p•. Therefore, we *de-novo* characterised repetitive elements in these last three species using RepeatModeler2 [91] and manually curated those sequences labelled as “LTR” and “Unknown” following the same method as in [92]. We also inspected consensus sequences with unusual classification for being avian repeats like many DNA transposon superfamilies [19]. We then concatenated the new *de-novo* libraries with the avian consensus sequences from Repbase [93], hooded crow [94], blue-capped cordon blue [95], flycatcher [96], and paradise crow [23], and used this final library to mask all six genomes with RepeatMasker [97]. The new repeat libraries are given in **Supplementary File S7**.

### Quantity of ERV transcription and their differential expression

We used Illumina RNA-seq reads from adult gonads from emu, chicken, and zebra finch (**Supplementary Table S1**) mapped against genomic copies/fragments annotated as LTR or ERV by RepeatMasker to quantify ERV transcription levels and investigate whether the ERVs were differentially expressed across available tissues. For these species, three to five biological replicates for every tissue were used.

Raw RNA-seq data were quality controlled using FastQC [98] and trimmed with TrimGalore [99] using default settings, then mapped to the respective reference genomes using STAR [100]. The alignment was filtered by running featureCounts function from the package Subread v2.0.0 in paired-end mode [101], and only uniquely mapping reads were retained. We provided featureCounts with a filtered RepeatMasker .out file containing only repeat copies annotated as LTR or ERV. Per-genome counts were obtained using read counts and lengths of corresponding ERVs. DESeq2 1.20.0 [102] implemented in the R Bioconductor package was used for relative quantification of the ERV transcripts and for calculating the TPM (transcripts per million), giving a normalized ERV expression level. Male reads that mapped to the W chromosome represented low counts and were therefore removed during the normalisation step. The values from replicas of each sample were averaged for the final plots of ERV expression. To identify biased ERVs per chromosome type (i.e., autosomes, Z, and W), we compared adult gonads from male and female individuals. The statistical analysis of differentially expressed ERVs was performed using DESeq2. All P-values were adjusted (*P*adj) using the Wald test. The degree of bias was determined by the log2 fold-change (Log2FC) difference between conditions. Therefore, the ERVs with log2FC > 0 and log2FC < 0 together with a *P*adj < 0.05 were considered as biased ERVs in the conditions.

### Full-length TE detection and abundance

Here we define full-length TEs as possible (retro)transposition-competent elements with relatively complete structures and the potential to produce transcripts. We identified fl-TEs in all the six avian genomes by adopting different methods for DNA transposons, LINEs (e.g., CR1), and LTR retrotransposons (ERVs). For DNA transposons and LINEs, we first identified open reading frames (ORFs) in the insertions annotated by RepeatMasker, then translated such ORFs and aligned with RPS-BLAST [103] against a custom Pfam [104] database containing transposon-related proteins (similar approach to [105]). ORFs from LINEs of at least 600 bp that spanned 90% of both endonuclease and reverse transcriptase domains were considered as full-length elements. Likewise, ORFs belonging to DNA transposons of at least 1 kb that spanned 90% of the transposase protein domain were considered full length.

In order to detect and quantify fl-ERVs, we used RetroTector [28] as well as LTRharvest [29] together with LTRdigest [30]. RetroTector results were filtered for score >300 and presence of 5’-LTR and 3’-LTR, as well as open reading frames (ORFs) with complete or partly complete *gag*, *pol*, and *env* genes as previously described in [28,106]. LTRharvest results were filtered for false positive using LTRdigest in combination with HMM profiles of LTR retrotransposon-related proteins downloaded from Pfam [104] and GyDB [107].

### Identification of SNVs of W-linked ERVs and their transcription and translation

To verify the hypothesis that the W chromosome is a refugium of intact and potentially active ERVs, we identified W-linked SNVs within ERVs and traced their transcription in RNA-seq data and translation in protein mass-spectrometry data wherever possible. W-linked ERV transcription was analysed in *Gallus gallus*, *Lycocorax pyrrhopterus* and *Taeniopygia guttata* (**Supplementary Table S1**). ERV translation was analysed in *Gallus gallus* white leghorn breed [37]. RNA-seq and proteome libraries selected for this analysis were from gonad tissue with the exception of *Lycocorax pyrrhopterus* for which the RNA-seq data was generated from female pectoral muscle.

To identify W-linked SNVs from male/female read mapping, we used the WhatGene pipeline developed by [108] for SNV analyses of B chromosomes and germline-restricted chromosomes [109] where we mapped male and female genome re-sequencing reads to the consensus sequences of our repeat library. We considered variants to be W-linked if they were present in all females but absent in males. We then checked for the presence of these W-linked variants in the RNA-seq data always following the WhatGene pipeline. Variants that were called W-linked from genomic data but were present in male transcriptomic data were discarded as false positives due to sample size.

To check for the presence of ERV-related proteins in white leghorn chicken proteome data, we extracted the ORFs from ERV consensus sequences and translated them into peptides using ORFFinder [110]. The peptide sequences were used as query database for MaxQuant 1.6.17.0 [38]. We used the experimental parameters described in [37] (**Supplementary File S5**); search results were filtered with a false discovery rate of 0.01. Second peptides, dependent peptides and match between runs parameters were enabled.

### Refugium index and toxicity index

To test whether intact TEs are uniformly distributed throughout the genome, we compared the observed total number of fl-ERVs (assuming that the numbers of other intact TEs are negligible in avian genomes [19]) on autosomes and sex chromosome to their expected values with a chi-square test with 2 degrees of freedom. We calculated the expected values of TE densities on the chromosomes by assuming a uniform density of these elements across chromosomes (**Supplementary Tables S3**-**4**). Next, we calculated the refugium and toxicity indexes, which are described below for SLCs in general.

The refugium index (formula **1.1**) calculates the percentage of excess or depletion of observed TE-derived bp (%TE_obs_) with respect to the genome-wide average of the total TE-derived bp of a haploid genome assembly (%TE_exp_). We recommend estimating TE densities in RepeatMasker or similar homology-based annotations using a species-specific repeat library combined with libraries of related species in Repbase or similar databases.

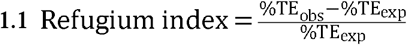

The refugium index indicates whether an SLC shows an excess (RI > 0) or a depletion of TEs (RI < 0). Furthermore, the refugium index can be estimated for any chromosome of interest, considering all TEs together or specific TE groups separately.

The toxicity index (formula **1.2**) calculates the excess of intact TEs present in the heterogametic sex with respect to the homogametic sex. Here 2n_hom_ and 2n_het_ are the total numbers of intact TEs in the diploid state in the homogametic sex (2 × autosomes + 2 × Z or X) and the heterogametic sex (2 × autosomes + 1 × Z or X + 1 × W or Y), respectively. We recommend quantifying intact TEs as the sum of the number of full-length LTR retrotransposons (incl. ERVs) in RetroTector/LTRharvest or similar structure-based approaches and the number of copies spanning >90% of the ORFs of DNA transposons (i.e., transposase) and LINEs (i.e., ORF1 or ORF2) in RPS-BLAST or similar homology-based searches.

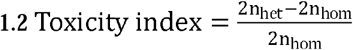

The toxicity index indicates whether there is no sex difference in toxicity (TI = 0), toxicity of the W or Y chromosome (TI > 0), or even toxicity of the Z or X chromosome (TI < 0). Consequently, we expect the toxicity index to be applicable not only to XY and ZW systems, but also XO systems.

## Acknowledgments

We thank Francisco Ruiz-Ruano for help with running WhatGene, Philipp Pottmeier for help with RNA extractions, Alexander J. Charles for help with running MaxQuant, and the Suh lab and Johannesson lab for helpful discussions. We thank Max Käller, Phil Ewels, Remi-André Olsen, Joel Gruselius, and Fanny Taborsak-Lines generated 10X data and assemblies at SciLifeLab Stockholm. We thank Erich Jarvis and the Vertebrate Genomes Project (VGP) for making their zebra finch, k•k•p•, and hummingbird assemblies available prior to publication, and Guojie Zhang and the B10K project for for doing the same with emu short read data. We thank Marco Ricci, Ivar Westberg, Jesper Boman, Diem Nguyen, and three anonymous reviewers for their comments on the manuscript. This research was supported by grants from the Swedish Research Council Formas (2017-01597 to AS; 2018-01008 to PJ), the Swedish Research Council Vetenskapsrådet (2016-05139 to AS; 2018-03017 to PJ; 621-2014-5113 and 2019-03900 to MI), the SciLifeLab Swedish Biodiversity Program (2015-R14 to AS), Villum Foundation (Young Investigator Programme, project no. 15560 to KAJ), and from the Carlsberg Foundation (Distinguished Associate Professor Fellowship, project no. CF17-0248 to KAJ). The Swedish Biodiversity Program has been made available by support from the Knut and Alice Wallenberg Foundation. Sequencing was performed by the SNP&SEQ Technology Platform in Uppsala, which is part of the National Genomics Infrastructure (NGI) Sweden and Science for Life Laboratory, and by the National Genomics Infrastructure in Stockholm. Both facilities are funded by Science for Life Laboratory, the Knut and Alice Wallenberg Foundation and the Swedish Research Council. Computations were performed on resources provided by the Swedish National Infrastructure for Computing (SNIC) through Uppsala Multidisciplinary Center for Advanced Computational Science (UPPMAX). We thank the State Ministry of Research and Technology (RISTEK); the Ministry of Forestry, Republic of Indonesia; the Research Center for Biology, Indonesian Institute of Sciences (RCB-LIPI); the Bogor Zoological Museum for providing permits to carry out fieldwork in Indonesia and to export select samples; the Natural Resources and Conservation Agency (BKSDA) Maluku, Ministry of Environment and Forestry-Republic of Indonesia. KAJ acknowledges a National Geographic Research and Exploration Grant (8853-10), the Dybron Hoffs Foundation and the Corrit Foundation for financial support for fieldwork in Indonesia.

## Data Accessibility

All the data is publicly available, and the code is accessible at http://github.com/ValentinaBOP/Wrefugium. All newly generated data were deposited in BioProject PRJNA604967 and Sequence Read Archive numbers are pending.

## Authors’ Contributions

VP and AS designed the study. VP analysed the data and wrote the first manuscript draft. VP and AS wrote and revised the subsequent drafts with input from all authors. KAJ and TH provided paradise crow samples. MI extracted paradise crow DNA. OMPG extracted paradise crow RNA and helped with the analysis of repeat transcription and their differential expression. JB helped with repeat annotation. PJ ran RetroTector analyses. QZ and JL provided the emu genome assembly. AS supervised the study. All authors read and approved the manuscript.

